# Diverse Mechanisms of *SMARCB1* Inactivation and Genome Maintenance Defects in Ultra-Rare Malignant Rhabdoid Tumors

**DOI:** 10.64898/2026.02.27.706889

**Authors:** Elizabeth Rasmussen, Elena Mironova, Zhao Lai, Kendra Maaß, Stefanie Volz, Dias Kurmashev, Stefan M. Pfister, Yidong Chen, Raushan T. Kurmasheva

## Abstract

Malignant rhabdoid tumors (MRTs) are extremely rare and highly aggressive pediatric cancers classically defined by biallelic loss of the *SMARCB1* gene, with rare involvement of *SMARCA4*. However, the molecular mechanisms leading to this loss are not yet fully understood. MRTs occur predominantly in infants, with the highest incidence in children under one year of age. Clinically, they are characterized by early metastatic dissemination and dismal outcomes, with 5-year event-free survival rates below 20%. There are currently no curative therapies for these patients.

Here, we performed integrated genomic, transcriptomic, and epigenomic profiling of 16 patient-derived MRT models including intracranial, renal, and soft tissue origins. While *SMARCB1* deficiency was ubiquitous, we observed substantial heterogeneity in the mechanisms driving its inactivation. Only two tumors harbored detectable coding single-nucleotide variants in *SMARCB1*; the predominant mechanisms involved large-scale deletions and broad loss-of-heterozygosity (LOH) on chromosome 22, with extensive LOH in tumors lacking point mutations or focal deletions, consistent with allelic loss as a frequent “second hit.” In contrast, *SMARCA4* remained intact across all models, reinforcing the mutual exclusivity of *SMARCB1* and *SMARCA4* alterations. Structural analyses revealed extensive variation, including more than 400 events per tumor on average and candidate gene fusions such as *AHI1:MYB*, whereas alterations in *TP53* and *BRCA1/2* genes were infrequent. Transcriptomic and epigenomic profiling showed heterogeneity driven by tissue of origin, disease progression, and therapeutic response, with subtype-specific programs and epigenetic modulation of DNA repair and immune-related genes (*SLFN11, MGMT, LIF*) linked to treatment sensitivity.

Collectively, our findings refine the molecular definition of MRTs, showing that while *SMARCB1* loss remains the foundational driver, tumor behavior is further shaped by structural variation, impaired DNA repair pathways, and dynamic epigenetic landscapes. These integrated changes contribute to tumor heterogeneity, progression, and differential therapeutic vulnerabilities. Beyond advancing mechanistic understanding and identifying candidate biomarkers for patient stratification, our multi-omics dataset represents a valuable resource for the research community, supporting future studies and efforts to improve clinical management of this highly aggressive pediatric malignancy.

## INTRODUCTION

Malignant rhabdoid tumors (MRTs) comprise a group of highly aggressive ultra-rare pediatric cancers. They consist of rhabdoid tumors of the kidney (RTK), atypical teratoid/rhabdoid tumors of the central nervous system (AT/RT), and extrarenal soft-tissue MRTs. These tumors are characterized by a high metastatic potential and poor prognosis, with a 5-year event-free survival rate of approximately 20%. Nearly all MRT cases harbor homozygous deletions of the *SMARCB1* gene (or rarely *SMARCA4*), a core subunit of the SWI/SNF (BAF) chromatin remodeling complex. This hallmark genetic alteration is observed in approximately 40% of extracranial MRTs, 25% of intracranial AT/RTs, and 70% of extrarenal MRTs^1–3^, and it underpins rhabdoid predisposition syndromes 1 and 2, the precise mechanisms of which driving MRT pathogenesis remain unknown^4,5^. Loss of *SMARCB1* follows a classic “second-hit” tumor suppressor model, in which an inactivating mutation is followed by loss of the remaining wild-type allele through mutation deletion, or loss-of-heterozygosity^6^. Despite their clinical aggressiveness, MRTs are genomically stable and largely diploid, with few recurrent amplifications or deletions^7–9^. The tumor-driving role of *SMARCB1* loss is believed to be primarily epigenetic, through dysregulation of chromatin remodeling and transcriptional programs, rather than through classic mutational oncogenic mechanisms. Nonetheless, therapeutic responses to agents targeting epigenetic regulators have been modest; for example, the EZH2 inhibitor EPZ011989 demonstrated limited efficacy in pediatric xenograft MRT models, as reported by the Pediatric Preclinical Testing Consortium (PPTC)^10,11^. Current treatment strategies, adapted from Wilms tumor protocols, include combinations of surgery, radiation therapy, and chemotherapy (e.g., vincristine, actinomycin, doxorubicin, with/without cyclophosphamide)^12,13^. Radiation therapy may be contraindicated in very young children, and while outcomes have improved in select subgroups, such as older children with localized, low-stage disease, overall survival remains limited, and curative targeted therapies are lacking^12,14^.

Emerging evidence supports the potential of PARP inhibitors as a novel therapeutic avenue for MRTs. Several PARP inhibitors are currently approved for various adult malignancies and are also being assessed in pediatric populations through phase I/II clinical trials, both as monotherapies and in combination regimens with DNA damaging agents. PARP inhibitors act by trapping PARP enzymes at DNA damage sites, thereby blocking repair and inducing cell death^15^. Talazoparib (TLZ), one of the most potent PARP1/2 inhibitors, demonstrates robust efficacy in *BRCA1/2*-deficient tumors and can be further potentiated in *BRCA1/2*-proficient models when combined with DNA-damaging agents^16–20^. Our prior work showed strong synergistic antitumor activity of TLZ combined with temozolomide (TMZ) in Ewing sarcoma models, and in a phase I/II Children’s Oncology Group trial, though toxicity limited dosing^21,22^. Next-generation, highly selective PARP1 inhibitors such as AZD-5305 are being clinically developed to improve therapeutic index and tolerability^23^. Additionally, novel nano-formulations, including PEGylated TLZ (PEG∼TLZ) and PLGA-encapsulated TLZ, enhanced tolerability while retaining potent anti-tumor efficacy in Ewing sarcoma models^16,17,24^. Notably, the combination of PEG∼TLZ and TMZ also elicited robust responses in MRT xenografts, independent of *SMARCB1* status^25^.

Although *SMARCB1* loss is the defining oncogenic event in MRTs, tumor cells retain DNA repair capability, suggesting that *SMARCB1* deletion drives tumorigenesis largely through epigenetic deregulation and signaling pathway perturbation. In our MRT models, we observed dysregulated pathways, including *EGFR*, *Ephrin*, and *FOXP1* signaling, associated with response to PEG∼TLZ and TMZ, and noted that MGMT (O^6^-methylguanine methyltransferase) protein expression inversely correlated with therapy response, highlighting its potential as a predictive biomarker^25–27^. Both PARP1 and MGMT are central to the repair of TMZ-induced DNA damage, providing a mechanistic basis for therapeutic synergy.

There remains a critical need to better understand the genomic and epigenomic landscape of MRTs to accelerate the development of effective therapies for this devastating disease affecting infants and children. To address this, we performed comprehensive multi-omic analyses of 16 MRT xenografts, representing intracranial (AT/RT-MYC and AT/RT-SHH), extracranial (RTK), and soft tissue subtypes. Whole genome analyses confirmed *SMARCB1* loss through diverse genomic mechanisms, accompanied by structural variation and loss-of-heterozygosity. Transcriptomic profiling revealed variability linked to tissue of origin, relapse, and therapeutic response, while genome-wide epigenetic characterization determined regulatory contributions to altered gene expression. Overall, our findings refine the molecular landscape of MRTs, uncover potential therapeutic vulnerabilities, and highlight the potential of PARP-targeted strategies in the pursuit of more effective and less toxic treatments.

## MATERIALS AND METHODS

### Nucleic acid isolation

KP-MRT-RY, KP-MRT-NS, KP-MRT-YM, and MP-MRT-AN xenograft tissues were a gift from Dr. Hajime Hosoi (Kyoto Prefectorial University). SJ-BT-12, BT-16, BT-29, BT-42, BT-55, WT-12, KT-14, WT-16, NCH-RBD1, NCH-RBD2, and Rh-18 xenografts were obtained from Dr. Peter Houghton (St. Jude Children’s Research Hospital and Nationwide Children’s Hospital). G401 was a cell line purchased from ATCC. Xenograft tumor tissues (**Table 1**) were ground under liquid nitrogen, and approximately 30–50 mg of the resulting powder was used for RNA and genomic DNA extraction and analysis. Genomic DNA was isolated using the Qiagen DNeasy Blood and Tissue Kit (*69506, Qiagen, Hilden, Germany*), following the manufacturer’s protocol for tissue samples. Briefly, the tissue powder was resuspended in 180 µL of ATL buffer and digested with 20 µL of Proteinase K at 56 °C for 1 hour, or until complete lysis was achieved. After digestion, 200 µL of AL buffer was added and mixed thoroughly by vortexing, followed by the addition of 200 µL of 96–100% ethanol. The mixture was then applied to a DNeasy Mini spin column, washed with 500 µL each of buffers AW1 and AW2, and the DNA was eluted in 50 µL of AE buffer. DNA concentration and purity were determined using a NanoPhotometer (*Model N50, Implen, Munich, Germany*).

**Table 1.**
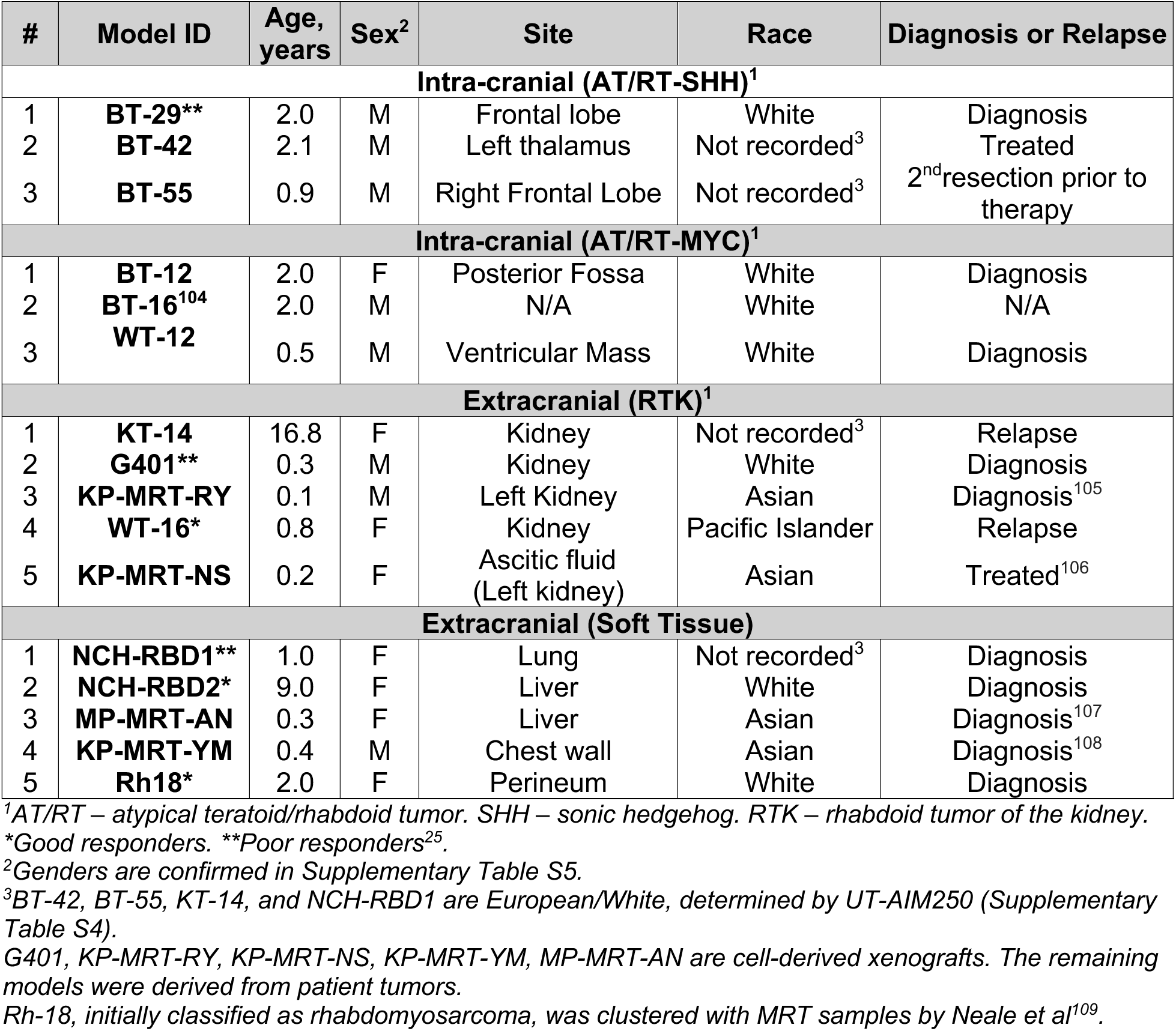
MRT xenograft models.

Total RNA was isolated from xenograft samples using the Qiagen RNeasy Mini Kit (*74104, Qiagen, Hilden, Germany*), according to the manufacturer’s tissue protocol. The tissue powder was homogenized in 600 µL of RLT buffer containing β-mercaptoethanol using a QIAshredder spin column (*79656, Qiagen*). The homogenate was centrifuged, and the resulting supernatant was combined with 600 µL of 70% ethanol and transferred to a RNeasy spin column. Bound RNA was washed using buffers RW1 and RPE and eluted in 50 µL of RNase-free water. RNA concentration and purity were also assessed using the NanoPhotometer (*Model N50, Implen, Munich, Germany*).

### DNA sequencing and variant analysis

Approximately 250–500 ng genomic DNA of 16 MRT samples was extracted and then the KAPA HyperPrep kit (*Roche Diagnostics, IN*) was utilized to construct DNA libraries for whole genome sequencing. Genomic DNA was sheared with a Covaris S220 Ultra Sonicator to the average of 200–400 bp fragments and then sequenced using 150 bp paired end sequencing at GCCRI Genome Sequencing Facility. For 16 MRTs, paired sequences were aligned to the hg38 genome using BWA aligner (v0.7.5a) and further processed following GATK’s (v4.3) best practice workflows (https://gatk.broadinstitute.org/). Two mutation callers, VarScan2 (v2.3.9)^28^ and GATK’s haplotypeCaller (v4.3) were used to determine variants from reference genome. Variants were further filtered by variant allele frequency (VAF) ≥ 0.1, a minimum depth of coverage of 20, minimum altered allele ≥ 3, and less than 1% of the altered allele in the population (1000 Genome Project). All the variants were annotated using ANNOVAR^29^. We considered variants, including SNVs (single-nucleotide variants) and indels, identified by both callers in our analyses. The combined Mutation Annotation Format (MAF) file from all tumors was generated using R/maftools (v.2.0.16)^30^, which is used for oncoPlots and other visualization plots. The structural variation (SVs) and indels from paired-end reads were called using MANTA caller (v.0.7.0)^31^ using the BAM files that were aligned to the hg38 human genome and passed through GATK pipeline. A panel of in-house curated ancestry informative markers (AIMs) from exonic regions^32^ were extracted and compared to the results from the 1000 Genome Project population background to determine the racial/ethnic background of all 16 samples.

### Identification of mutational signatures

The MAF file converted from 16 VCFs were processed with maftools as previously described. The 96 classes of single base substitution (SBS) mutation signatures were accessed from the publication by Alexandrov et al ^33^. A signature discovery software R/MutationalPatterns (v3.8.1) were used bootstrapping method with 1000 iterations to avoid signature misattribution and for the estimation of the percentage and mean contributions of all signatures^34^.

### Loss-of-heterozygosity (LOH)

Single-nucleotide polymorphisms (SNPs) for genome-wide LOH assessment were derived from Affymetrix Genome-Wide Human SNP Array 6.0 (NCBI/GEO GPL6801, annotated in UCSC hg19). A total of 929,727 SNPs were selected and converted to UCSC hg38 annotation (UCSC Genome Browser, LiftOver tool. Total of 929,324 SNPs passed through the LiftOver). Variant fractions and coverage depths were obtained using varScan2, which were used to assess DNA gain or loss from coverage depth, and LOH from allelic fraction. Visualizations of copy number variation (CNV) and LOH were done by using Broad Institute’s Integrated Genome Browser (IGV)^35^ and R/karyoploteR (v1.10.5)^36^.

### Gender confirmation

Since germline DNA data were unavailable, sex was determined for 16 MRT samples using RNA-seq data, primarily by examining expression of two marker genes: *XIST*, a long noncoding RNA involved in X-chromosome inactivation in females, and *RPS4Y1*, a Y-chromosome-encoded gene expressed almost exclusively in males.

### Protein extraction and Western blotting

Xenograft tumor tissues were ground under liquid nitrogen and ∼30-50 mg of powder was used for Western blot analysis. For protein extraction, tissue powder was lysed with ice-cold RIPA lysis buffer supplemented with protease and phosphatase inhibitors. After ultra-sonication, the lysates were centrifuged at 12,000 rcf (relative centrifugal force) for 20 min at 4°C. Lysate protein concentrations were determined using the BCA protein array kit (*5000002EDU, Bio-Rad*). Equal amounts of protein from each sample were mixed with SDS loading buffer (*NP0007, Invitrogen*), separated on 10% Bis-Tris gel (*Invitrogen™ NuPAGE™, NP0302BOX*), and transferred to Immobilon®-FL PVDF Membrane (*Millipore, IPFL00005*) using an iBlot transfer system. The membranes were probed with antibodies against SMARCB1 (*1:1000, Cell Signaling, 91735S*) and GAPDH (1:1000, *Cell Signaling, 2118*). Proteins were detected with LI-COR IRDye 680RD anti-mouse and LI-COR IRDye 800CW anti-rabbit secondary antibodies (1:20,000) in the mixture of LI-COR Odyssey® Blocking Buffer (TBS), 0.2% Tween 20, and 0.01% SDS. Membranes were imaged using LI-COR Odyssey® CLx Imaging System (*Nebraska, USA*). The intensity of each protein band was quantified using Li-COR Image Studio software and normalized against the corresponding GAPDH signal. The normalized values were used to plots (Prism 9 GraphPad).

### RNA sequencing and differential expression (DE) analysis

RNAseq libraries were prepared for each sample by following the NEB Directional mRNA-seq sample preparation guide (*New England Biolabs, Ipswich, MA*). The first step in the workflow involved purifying the poly-A containing mRNA molecules using poly-T oligo-attached magnetic beads. Following purification, the mRNA was fragmented into small pieces using divalent cautions under elevated temperature. The cleaved RNA fragments were copied into first strand cDNA using reverse transcriptase and random primers. These cDNA fragments then went through an end repair process, the addition of a single ‘A’ base, and then ligation of the adapters. The products were then purified and enriched with PCR to create the final RNA-seq library. RNA-seq libraries were subjected to quantification and subsequent 100bp paired read sequencing run with Illumina NovaSeq 6000 platform (Greehey Children’s Cancer Research Institute (GCCRI) Genome Sequencing Facility (GSF) at UTHSCSA).

Upon obtaining the sequence reads (∼130–200 million reads, or 75–100 million pairs), the samples were aligned to USCS hg38 human genome and mm10 mouse genome using HISAT aligner (v2.1.0)^37^. We assigned each sequence read to 4 groups, human-only, mouse-only, common, and unaligned, by using the following criteria: (1) alignment scores, (2) number of mismatched bases, and (3) length of matched segment length, from paired-end reads. To remove mouse RNA contamination, we extracted all reads from human-only and common groups (∼78–94% reads) and then aligned to the human genome (hg38) using HISAT followed with Stringtie (v2.1.3b) to quantify gene expression levels (both in read counts and in Fragment Per Kilobase of transcript (FPKM) for all RefSeq genes^37^. Differential expression (DE) analyses were performed using DESeq v1.38.0 (R/Bioconductor) for the various tissue-tissue comparisons, “good responders” versus “poor responders”, and “relapsed” versus tumor collected at time of diagnosis (“diagnosis”) groups^38^. Up- and downregulated genes were determined by the following criteria: (1) absolute log2 fold change (log2-FC) > 1, (2) average FPKM > 1, and (3) multiple-test-adjusted (Benjamini–Hochberg) *p* value < 0.05. Using the differential expression fold change, we also performed Gene Set Enrichment Analysis (GSEA)^39^, using the stand-alone package from the Broad Institute (v4.0.3) or the implementation provided by the ClusterProfiler package v3.12.0 (*R/Bioconductor*)^40^. Additional functional analysis were performed using Ingenuity Pathway Analysis (*IPA,* https://digitalinsights.qiagen.com/*, Qiagen*), and STRING (https://string-db.org/)^41^ for protein-protein interaction network and biological function enrichment.

### Gene fusion identification and one-step RT-PCR analysis

The gene fusion detection/analysis was performed using STAR-Fusion (v1.1.0) (https://github.com/STAR-Fusion) algorithm using hg19 human genome. The --min_junction_reads number was set to 5 and the rest of parameters were set to defaults. The fusion results were further processed using R to constrain the spanning reads to at least 5 and using R/chimeraviz (v1.34.0) package for circle plot and fusion plot gene plots.

To validate the *AHI1:MYB* fusion transcript identified in KP-MRT-NS, we designed primers flanking the predicted fusion junction (forward: 5′-CTTGCCATTCAAGTCACCCA-3′ and reverse: 5′-TGTCTCACATGACCAGCGTC-3′), along with control primers targeting *AHI1* (forward: 5′-AGGAGCAGCAAATTATCGGGA-3′ and reverse: 5′-TGTCTCACATGACCAGCGTC-3′). The expected amplicon sizes were 237 bp for the fusion product and 159 bp for the *AHI1* control. Total RNA was isolated from 30 mg of snap-frozen patient-derived xenograft (PDX) tissue using the RNeasy Mini Kit (*74004, Qiagen*). One-step RT-PCR was performed using the Invitrogen One-Step RT-PCR Kit (*12596025, ThermoFisher Scientific*), to amplify the predicted fusion junction from the same PDX sample used for sequencing (KP-MRT-NS). As a positive control for *AHI1* expression, we used PDX sample UTSW-2264 (Wilms tumor), which expresses high levels of *AHI1* mRNA transcript (TPM = 70.77)^42^. Each RT-PCR reaction contained 50 ng of total RNA in a 50 μl reaction volume, with all steps carried out in accordance with the manufacturer’s protocol. Thermal cycling conditions were: 50°C for 30 min, 95°C for 2 min, followed by 40 cycles of 95°C for 15 s, 55°C for 30 s, and 65°C for 1 min. RT-PCR products were resolved on a 2% agarose gel and visualized using GelRed Nucleic Acid Stain (*41003, Biotium*) on a BioRad imaging system (*ChemiDoc MP*).

### Enzymatic methylation sequencing and data pre-processing

Sixteen rhabdoid tumors’ DNA were processed using NEBNext Enzymatic Methylation Kit (EM-seq) for Illumina library construction and accurate representation of 5mC within the tumor genome. Briefly, the first enzymatic step involves the oxidation of 5-methylcytosine to 5-carboxycytosine using TET2 and the glycosylation of 5-hydroxymethylcytosine using T4-BGT, and then the DNA is made single-stranded using either formamide or sodium hydroxide according to the manufacturer’s protocol. APOBEC then deaminates cytosines into uracils. Since 5-methylcytosines and 5-hydroxymethylcytosines were protected in the first enzymatic step, these forms are no longer substrates for APOBEC and are not deaminated. Illumina libraries were constructed by following the NEB EM-seq kit protocol^43^.

FASTQ data derived from EM-seq protocols of 16 MRTs were piped into FastQC (v0.12.1) to check data quality. Reads were trimmed using Fastp (v0.23.2) with Illumina’s standard adapters^44^. BWA_meth (v0.2.2) was used to align reads to human genome (UCSC hg38 build) using default parameters. SAMtools (v1.19.2) was used to further convert the data to BAM format for MethylDackel (v0.3.0) to process into bedGraph format that contains chromosomal position, methylation percentage, number of reads reporting methylated bases, and reads reporting unmethylated bases^45,46^. We used CpG island definitions from Illumina’s MethylationEPIC v2.0 manifest (from the “Relation_to_UCSC_CpG_Island” column) and their associated gene names from the “UCSC_RefGene_Name” column. Python package CpGtools (v2.0.2, on Python v3.9.18) was used (CpG_aggregation.py) to isolate all CpGs islands within promoters from the bedGraph files per tumor with the parsed EPICv2 CpG islands as input^47^. Each CpG island was further annotated (CpG_anno_position.py) with its associated gene names. The *beta (β) value* of each CpG island was calculated separately in base R as *β* = *C*/(*C* + *U*) where *C* is the methylated reads and *U* is unmethylated reads. Principal Component Analysis (PCA) were performed (beta_PCA.py) to represent the relationships between tumors (both unsupervised and supervised). All CpG islands identified as differentially expressed in the volcano plot input files of the tissue-tissue comparisons were aggregated into a single file for supervised PCA plot generation.

To facilitate the differential methylation analysis, we also estimate *coverage* for each tumor per isolated CpG island using SAMtools bedcov (per base read depth) and then divided by the length of the read to produce the average read count. Average coverage per tumor group was also calculated. R/methylKit (v4.5.1)^48^ was used to produce percent methylation and coverage histograms per tumor for quality control purposes.

### Differential methylation (DM) of promoter CpG islands

Differential methylation calculations by promoter CpG island were performed using CpGtools (dmc_fisher.py) for all group comparisons (“AT/RT vs RTK”, “AT/RT vs Soft Tissue”, “RTK vs Soft Tissue”, “Diagnosis vs Relapse”, and our previously published “Good vs Poor Responders” groups^25^), from which log2 Odds Ratio 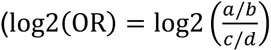 where ‘a’ is the sum of all methylated reads of tumors in the first group, ‘b’ is the sum of all unmethylated reads of tumors in the first group, ‘c’ is the sum of all methylated reads of tumors in the second group, and ‘d’ is the sum of all unmethylated reads of tumors in the second group), p-value (Fisher’s exact test) and Benjamini-Hochberg adjusted p-value for each CpG Islands were evaluated for DM. After initial processing (removing duplicated CpG entries and CpG entry rows containing NAs), significant differentially methylated promoter CpG islands were selected based on following criteria: (1) multiple test-adjusted p-value > 0.01; (2) absolute(log2-OR) > 4, and (3) average coverage per feature in both groups > 10 (at least 10x sequencing coverage). In addition, we further constrained the consistency with an unpaired two-tailed Student’s t-test p-value less than 0.2 or 0.25 (depending on the comparison, or half the rounded mean of the comparison’s t-test p-values per differential methylation analysis). The additional step will further remove cases with extreme value at one sample only. Volcano plots for DM islands were generated using R/ggplot2 (v3.5.2) with hypo- and hyper-methylation marked in blue and red color, respectively. The top 20 hypo- and hyper-methylated genes (based on adjusted p-value) were labelled to assist the visualization. DM genes were combined with differentially expressed (DE) genes (derived from DESeq analysis) to visualize the regulation relationship between DM and DE genes. Genes that were significant in DM, DE, significant in both DM and DE (labeled), or neither, were plotted in red, blue, green and gray, respectively. Due to the nature of CpG islands, we only focus on second and forth quadrants for possible inverse regulation relationship in the final log2-OR and log2-FC quadrant plots for each comparison.

### Intracranial AT/RT sub-classification

Methylation predictions for six EM-seq samples were generated using the MPACT classifier. To compare EM-seq-based methylation profiles with an established reference dataset, we used methylation data from the central nervous system (CNS) tumor classification study by Capper et al., which forms the basis of the Molecular Neuropathology (MNP) classifier ^49^. From this reference cohort, only the AT/RT samples representing the subgroups AT/RT-SHH, AT/RT-MYC, and AT/RT-TYR were selected.

Raw IDAT files from the reference cohort samples were processed using the minfi package to obtain CpG-level beta values, which were combined into a unified CpG-by-sample matrix^50^. For the EM-seq dataset, all six bedGraph files (containing methylation scores ranging from 0 to 100) were imported using rtracklayer^51^, coverage fields were removed, and methylation scores were scaled to beta values between 0 and 1. Genomic coordinates for all EPIC CpGs were obtained from the EPIC hg38 manifest and converted into genomic ranges^52^. EM-seq methylation values were assigned to EPIC CpGs based on genomic overlap, and CpGs without overlap were recorded as missing. The resulting EM-seq beta matrix and the Illumina array beta matrix were merged using shared CpG identifiers to ensure genomic alignment across platforms.

To visualize how EM-seq samples relate to the known AT/RT methylation subgroups, we performed t-SNE with the Rtsne package on the harmonized dataset using the top 10,000 most variable CpGs^53^. The two-dimensional embedding was visualized using ggplot2 together with the subgroup labels defined in the reference dataset^54^.

### Statistical analyses

Data were statistically analyzed and presented as the mean ± SEM (standard error of the mean). Unless otherwise noted, statistical significance (p-value) was determined by a not-paired two-tailed Student’s t-test using Excel (v16.78 (23100802) and Prism (v10.2.3(347)) software. For multiple group comparison, we used the one-way ANOVA, followed by Tukey’s Honest Significant Difference (HSD) method (R, http://www.R-project.org). For linear regression with scatter plots, we report the 95% confidence interval, Pearson correlation coefficient, and corresponding p-value using the R/ggpubr package (v0.4.0).

## RESULTS

### *SMARCB1* Deletion Status Across Tumors

We performed WGS on 16 MRT tumors, comprising 6 AT/RT, 5 RTK, and 5 soft tissue (**Table 1**). Sequencing yielded an average of 483 million reads per sample aligned to the human reference genome (hg38) at a mean depth of ∼67-fold (150 bp paired-end reads; **Supplementary Table S1**). After applying variant-calling filters requiring: (1) ≥ 20-fold coverage, (2) ≥ 20% variant allele fraction with ≥ 3 supporting alternate reads, and (3) <1% population frequency in the Exome Aggregation Consortium (ExAC) ^55^, we identified a total of 8,187 single-nucleotide variants (SNVs, ∼512 per tumor) and 524 small insertions/deletions (indels) across the cohort (∼33 per tumor, Supplementary Table S2).

To examine the DNA alterations at SNV or indel levels in genes previously associated with MRT, we analyzed a curated panel of 27 genes (**Figure 1A**)^7,56,57^. *TP53* alterations were detected in 11 of 16 tumors (69%), all carrying the P72R variant (**Supplementary Table S3**); with only KP-MRT-NS harboring an additional R273C mutation (**Figure 1B)** ^58^. The P72R variant retains partial *TP53* tumor suppressor function^59^, and represents a common, widely distributed polymorphism that has been associated with increased cancer risk in certain populations^60–63^. The R273C variant is a recurrent oncogenic hotspot associated with many cancers and poor prognosis ^64^. To focus on potentially relevant variants, we further filtered *TP53* and *BRCA1/2* mutations to retain only those with a general population frequency below 1%. Notably, 3 tumors with *BRCA1* mutations also carried *TP53* alterations, and one of these (WT-16) had an additional *BRCA2* mutation (**Figure 1A-B, Supplementary Table S3**). Only 2 tumors, WT-12 (p.R201X) and KP-MRT-RY (p.R53X), contained previously reported *SMARCB1* mutations, with p.R201X located within the SNF5 region of the SWI/SNF chromatin-remodeling complex^65^. Mutation counts in the remaining genes were low (one variant each in *TP53BP1*, *PARP1*, *PARP3*, *SPECC1L*, *TTC28*, and *SUZ12,* and none in the others). No *SMARCA4* mutations were detected in any sample. WT-16 demonstrated the highest overall mutation burden (1,468 SNV and indels), including mutations in *TP53, BRCA1, and BRCA2*.

**Figure 1.**
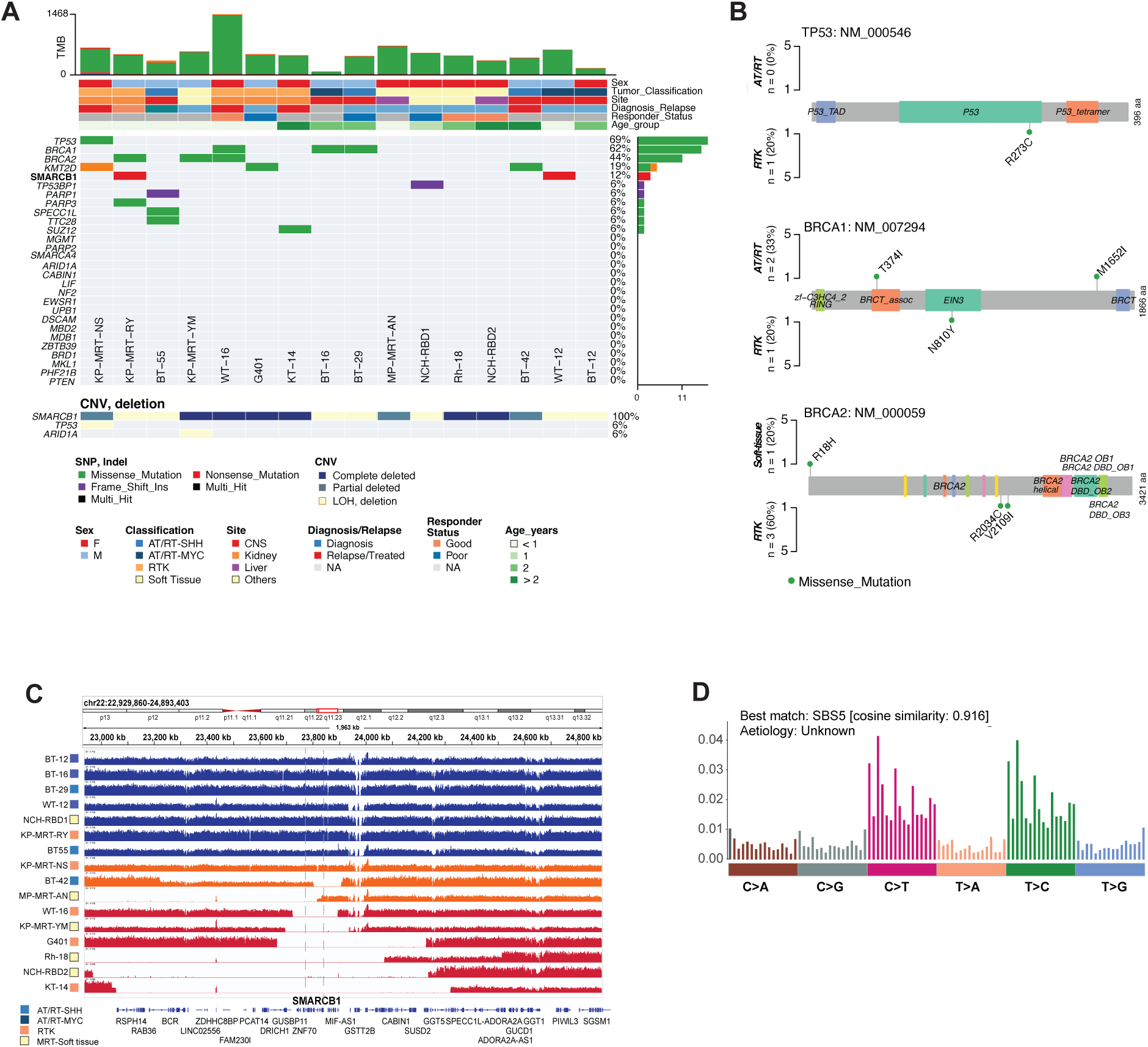
Mutation spectrum (SNPs and INDELs) across 16 MRT models. (**A**) Mutation and copy number profiles from a selected panel of genes implicated in MRT. The top panel contains tumor mutation burden (TMB, unit of mutation counts across genome), and key demographic/clinical parameters. The left panel illustrated the percent of MRT models carrying one or more mutations. The CNV panel shows gene-level copy number alternation, including loss-of-heterozygosity. (**B**) Gene-level mutations detected in *TP53, BRCA1* and *BRCA2*. Mutations were selected based on the following criteria: (1) sequence coverage >20x, (2) Altered allele t observed ≥3 times, (3) variant allele frequency >20%, and (4) variant allele in population frequency <1%. For *TP53*, the ten samples carrying the P72R variant were omitted from the lollipop plot. Similarly, *BRCA1* and *BRCA2* variants with a population frequency greater than 1% were excluded. (**C**) Deletion status of the *SMARCB1* region. Samples are color-coded as red (complete deletion), orange (partial deletion), and blue (“normal”). (**D**) Mutational signature analysis identified a single signature resembling SBS5 (cosine similarity = 0.916), determined using Non-negative Matrix Factorization (NMF) method.

Given that only 2 tumors carried SNVs in *SMARCB1*, we closely examined alterations on chromosome 22, where *SMARCB1* resides. Six samples (WT-16, KP-MRT-YM, G401, Rh-18, NCH-RBD2, and KT-14) showed large-scale deletions encompassing the *SMARCB1* region (**Figure 1C, red tracks**), while 3 samples (KP-MRT-NS, BT-42, and MP-MRT-AN) exhibited partial deletions (**Figure 1C, orange tracks**). Mutational signature analysis identified a single detectable signature, SBS5 (cosine similarity = 0.92), whose proposed etiology includes defects in nucleotide excision repair^66^, consistent with prior MRT studies^7^ (**Figure 1D**).

We further inferred genetic ancestry for all 16 MRT samples using a panel of 250 ancestry-informative markers located in exonic regions and derived from reference populations in the 1,000 Genome Project ^67^, following methods described in ^32^. Estimated ancestry proportions for each sample are summarized in **Supplementary Table S4** and corroborate the annotations provided in **Table 1**. We also performed gender analysis using RNA-seq data, with the results summarized in **Supplementary Table S5**.

### *SMARCB1* Loss-of-Heterozygosity (LOH) and Other Genome-Wide Structural Variations

We next performed genome-wide copy-number alteration (CNA) analysis across all 16 MRT samples (**Figure 2A**). We observed that NCH-RBD-1, KP-MRT-NS, and KP-MRT-RY displayed the most extensive genomic disruptions. Chromosomes 1, 2, 5, 7, 11, 12, and 17 each contained alterations (gains and losses) in at least 2 samples. We examined the *SMARCB1* locus on chromosome 22 in greater detail (**Figure 2B, left panel**). Importantly, all samples in the blue track, which lack *SMARCB1* SNVs or detectable deletions, showed broad loss of heterozygosity (LOH) across the region (**Figure 1A**). This pattern likely represents an additional mechanism of *SMARCB1* loss of function. Similar analyses were performed for *SMARCA4*, *TP53* and *ARID1A*.

**Figure 2.**
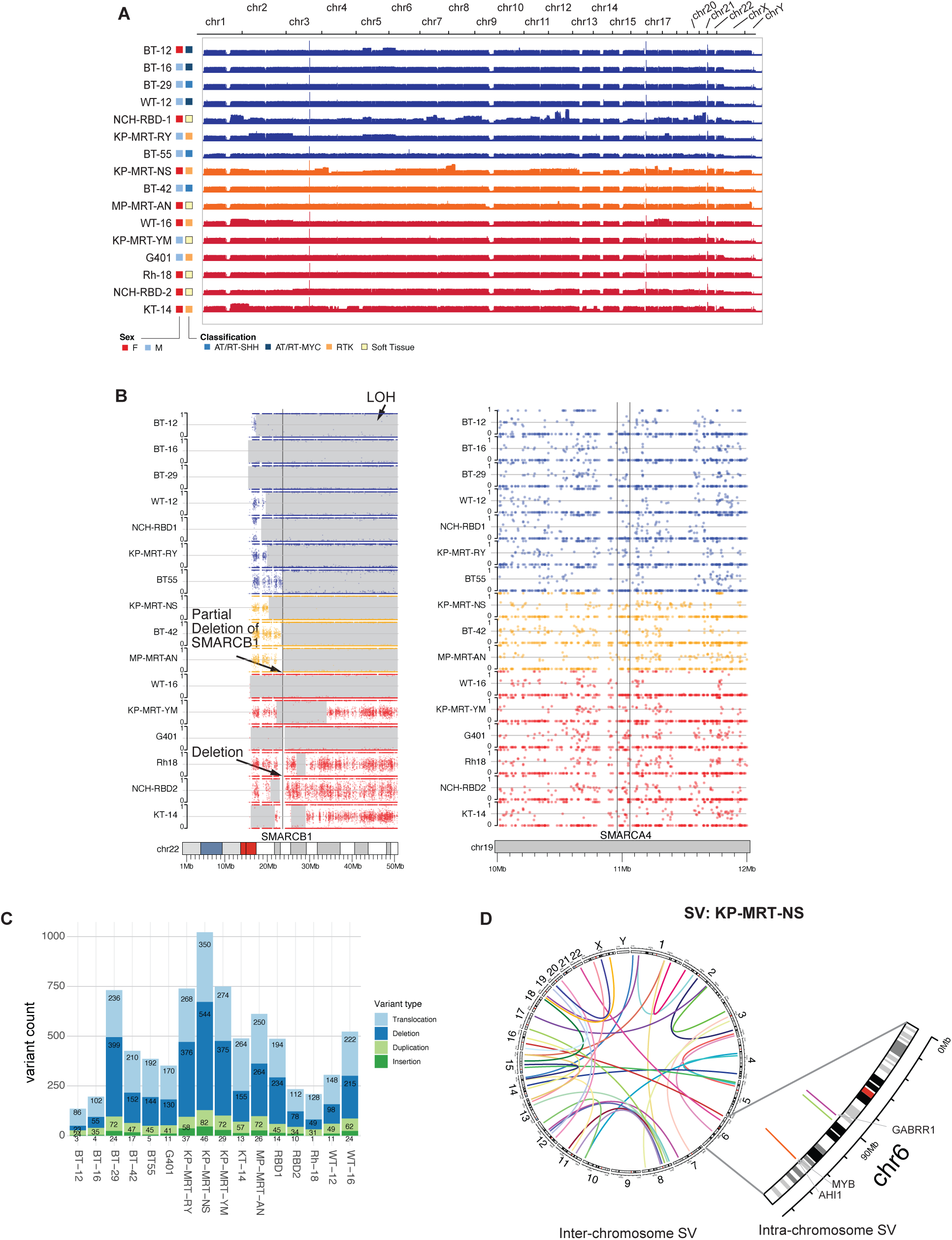
Genome-wide gain/loss, LOH, and structural rearrangements. (**A**) Genome-wide DNA alteration landscape of 16 MRTs are generally quiet, with except of NCH-RBD1 and KP-MRT-RY. Tracks are color-coded according to *SMARCB1* CNV status (Figure 1C). (**B**) Loss-of-heterozygosity (LOH) statuses of *SMARCB1* and *SMARCA4*. LOHs were assessed using a panel of ∼1 million SNPs (see Methods) across the genome. Each dot represents the observed allele fraction. Grayed regions indicate LOH, where one allele was detected or both alleles are identical (allele fractions clustered at ∼0 or ∼100%). No LOH was observed in the *SMARCA4* region of 16 MRT models. (**C**) Structural variations (SV) detected in each model, including translocations (BND, or breakend), deletions (DEL), duplications (DUP), and insertion (INS). (**D**) A Circos plot shows all inter-chromosomal translocations (n = 47) of sample KP-MRT-NS (with at least translocation observed at least 5 times and >10% fraction in both split- or spanning-reads), intra-chromosomal translocation (n = 96), highlighting a duplicated regions on chromosome 6 near *MYB* and *AHI1* loci.

*SMARCA4* remained intact in all tumors, with no evidence of SNVs, indels, or LOH. LOH events were identified in only one sample for each *ARID1A* (KP-MRT-YM) and *TP53* (KP-MRT-NS).

We next assessed structural variation (SV) across the genome using the MANTA algorithm^31^, focusing particularly on translocations annotated as breakends (BNDs). As summarized in **Figure 2C**, using the selection criteria described in the Methods, tumors harbored an average of 475 SV calls per tumor, including approximately 200 rearrangements, 206 deletions, 52 duplications, and 17 insertions (**Supplementary Table S6**). Interestingly, within KP-MRT-NS, we also identified one region covering *AHI1* (*Abelson Helper Integration Site 1*) and *MYB* (*MYB Proto-Oncogene*), where two distinct copy-number gains were detected (**Figure 2D**). In contrast, KP-MRT-RY sample has similar number of inter-chromosomal SVs, but different intra-chromosomal SVs (non-recurring comparing to KP-MRT-NS) as shown in **Supplementary Figure S2**. These local duplications do not span the full coding regions of either gene, and additional transcriptomic analyses will be required to determine their functional significance.

### Transcriptome Variability

We first evaluated *SMARCB1* expression by Western blot analysis (**Figure 3A; Supplementary Figure S1**). Among the 16 MRT samples, only NCH-RBD-1 demonstrated a strong signal, consistent with our earlier report analyzing 6 MRTs^25^. To investigate whether transcriptomic variation across different tumor statuses (tissue of origin, diagnosis/relapse, and treatment response), we performed whole-transcriptome analyses. We initially conducted differential expression (DE) analysis comparing 3 MRT subtypes: AT/RT (n = 6), kidney (n = 4), and soft-tissue (n = 6), using a threshold of ≥ 2-fold change and p-value < 0.05. This analysis identified 268 differentially expressed genes (DEGs) that clustered into 4 distinct expression patterns (**Figure 3B**). Functional enrichment analysis revealed subtype-specific pathways: interferon alpha/gamma responses in AT/RT, xenobiotic metabolism in kidney tumors, and epithelial mesenchymal transition (EMT) in soft-tissue tumors. Using the same criteria, we compared samples collected at initial diagnosis (n = 11) versus relapse or post-treatment (n = 4), identifying 68 DEGs (**Figure 3C**). Notably, genes upregulated in relapse samples included *EFS* (*Embryonal Fyn-associated substrate*, log2FC = 6.9, p < 0.01) and *FGF19* (*Fibroblast Growth Factor 19*, log2FC = 9.87, p = 0.01), while down-regulated genes included *S100A11* (*Calcium Binding Protein in the S100 family*, log2FC = −6.01, p < 0.01) and *PRKCDBP* (also known as *CAVIN3 caveolae associated protein 3*, log2FC = −9.50, p < 0.01).

**Figure 3.**
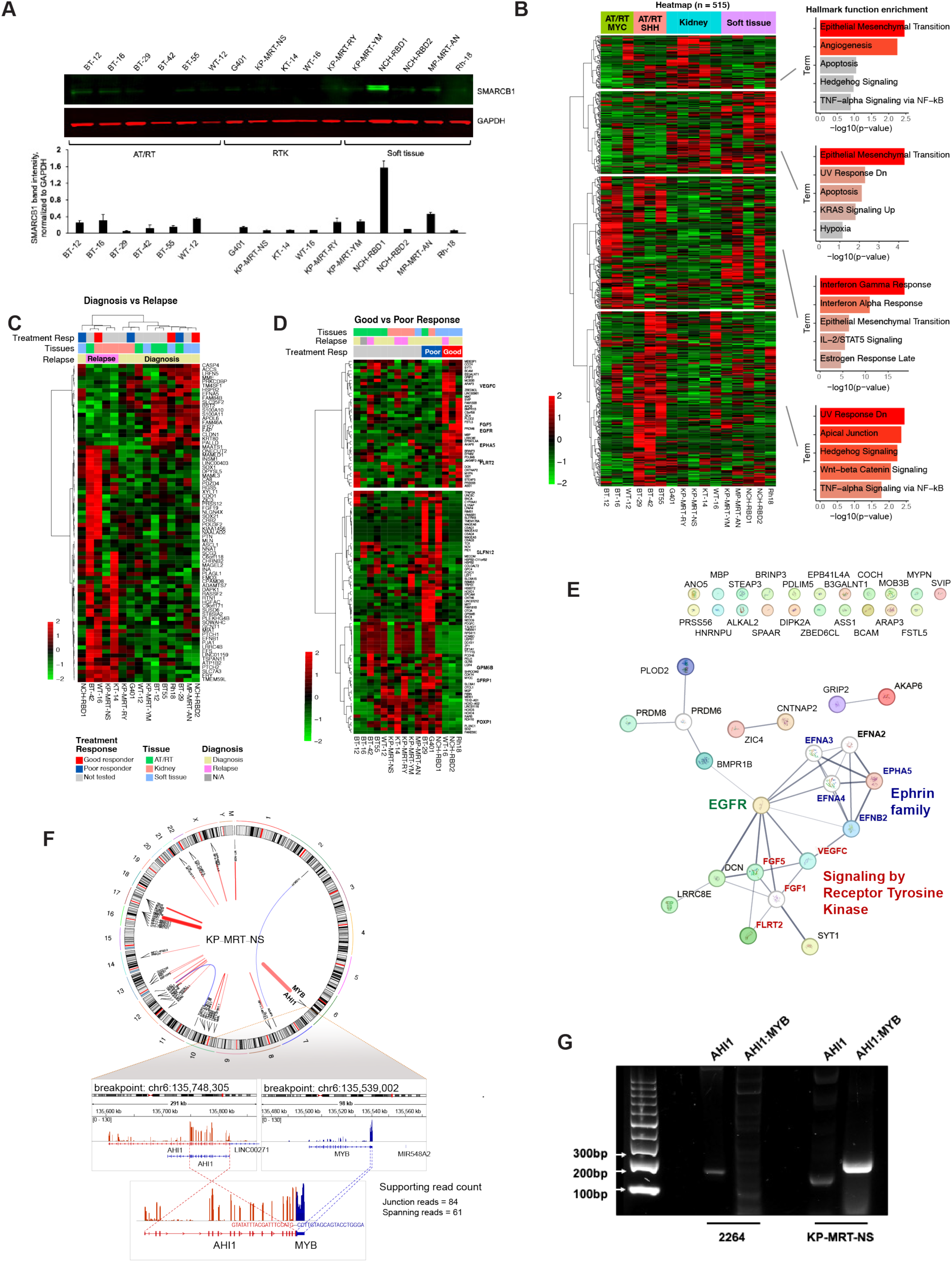
Transcriptomic changes in MRT models. (**A**) Consistent with the DNA-level SMARCB1 alteration shown in Figures 2A and 2B, Western blot analysis confirms loss or reduction of SMARCB1 protein (46 kDa) expression across all 16 MRT samples except for NCH-RBD1. GAPDH (37 kDa) served as the loading control (top). Quantification of band intensities normalized to GAPDH loading control was plotted (bottom). (**B**) Heatmap of differentially expressed genes derived from comparing AT/RT to RTK. Four unique transcriptional programming were identified, and their corresponding top 5 Hallmark gene sets (mSigDB) enrichment were plotted (the full lists of DEGs are provided in **Supplementary Tables S20-24**). (**C**) Heatmap of differentially expressed genes derived from comparing tissues obtained at relapse (or treated) samples obtained at diagnosis (the full lists of DEGs are provided in **Supplementary Table S25**). (**D**) Heatmap of differentially expressed genes comparing three responders to PEGylated talazoparib (PEG∼TLZ), temozolomide (TMZ), or combination therapy with three non-responders (n = 3 for each group), and the full lists of DEGs are provided in **Supplementary Table S26**). (**E**) Functional enrichment analysis (STRING) of the top 40 upregulated DEGs in responders revealed enrichment in EGFR signaling, receptor tyrosine kinases, and Ephrin family genes. (**F**) Example of fusions identified using STAR-fusion algorithm in sample KP-MRT-NS, visualized using a Circos plot. For *AHI1-MYB* fusion (same region as in Figure 2D), their expression levels, and the corresponding fusion product are shown (*AHI:* flipped, red; *MYB*: blue). (**G**) Primers spanning the fusion breakpoint and the 5′ region of *AHI1* (positive control) were designed, and RT-PCR validation results are provided. On a 2% agarose gel, the UTSW-2246 control sample (Wilms tumor PDX) showed only the 159 bp *AHI1* control band, with no detectable fusion product, confirming assay specificity. In contrast, the KP-MRT-NS sample exhibited both the 237 bp *AHI1:MYB* fusion band and the 159 bp *AHI1* control band, confirming the presence and expression of the fusion transcript in this tumor sample.

Among the 16 xenograft models, 6 were previously evaluated for anti-tumor activity of PEGylated talazoparib (PEG∼TLZ), temozolomide (TMZ), and their combination^25^. Tumor volumes were measured weekly for up to 13 weeks and objective tumor responses were categorized as partial response (PR) (NCH-RBD-1, BT-29) or progressive disease (PD) (G401) as poor responders, and complete response (CR) (Rh-18) or maintained complete response (MCR) (NCH-RBD-2, WT-16) as good responders (**Figure 3D**). Differential gene expression analysis comparing good versus poor responders yielded 116 DEGs (p < 0.01 and > 2-fold change). Functional analysis using STRING with 40 up-regulated genes (allowing connections to second shell with <5 interactors) revealed enrichment in Ephrin family: *EFNA2-5* and *EFNB2* (log2-fold change of −3.42, −1.38, −1.71, −0.95, 3.70, respectively), Receptor Tyrosine Kinase (RTK) family: *VGFC, FGF1/5, FLRT2* (log2FC = 8.70, 1.74, 7.69, 3.12, respectively), and epidermal growth factor receptor: *EGFR* (log2-FC = 5.41). (**Figure 3E**; see **Supplementary Figure S2** for downregulated genes). Notably, high *EGFR* expression may indicate elevated DNA damage^68–70^, suggesting that tumors with defective DNA repair pathways could be more susceptible to PARP inhibitors, such as TLZ. Two additional MRT models (KP-MRT-NS and MP-MRT-AN) also showed increased *EGFR* expression, similar to that observed in models sensitive to TLZ, TMZ, or their combination. We further examined whether intronic mutation impacts gene expression by grouping samples according to intronic mutation status for each gene. Several genes, including *CABIN1* and *BCR*, showed significant differential expression (**Supplementary Figure S3,** p = 2.3 x 10^-6^ and 4.02 x 10^-4^, respectively). Notably, both genes are in close genomic proximity to *SMARCB1*.

From paired-end RNA sequencing of 16 MRT models, we detected 268 fusion transcripts with the STAR-Fusion algorithm, requiring a minimum of 5 junction reads. After filtering for fusions with fewer than 5 spanning reads, 59 fusions remained (41 unique to a single sample, 18 found in ≥ 2 samples), including *ERBB4:IKZF2*, *SPECC1L:PI4KA*, and *TMEM123:YAP1*. Candidate fusions were further cross-referenced with the FusionGDB2 database (**Supplementary Table S7**). Only 17 unique fusion events were annotated in FusionGDB2, indicating these fusions have been reported in prior studies, whereas the remaining events are likely to be either sequencing or alignment artifacts ^71^. Among these, the *AHI1:MYB* fusion (left breakpoint: AHI1:chr6:135748305:-, right breakpoint: MYB:chr6:135539002:+) identified in sample KP-MRT-NS (**Figure 3E**) was selected for validation based on its high number of spanning and junction reads. The *TMEM123:YAP1* fusion also showed similarly high read report. RT-PCR, using the primers listed in **Supplementary Table S7** confirmed expression of this fusion transcript (**Figure 3F-G**).

### Genome-Wide Epigenetic Alterations

Genome-wide DNA methylation profiling was performed on 16 MRT models using EM-seq protocols. To assess overall epigenetic alterations, we first aggregated all promoter CpG islands (n = 15,507 of 25,238) and applied unsupervised principal component analysis (PCA) in 2-dimensional space (**Figure 4A**). Soft-tissue tumors and RTKs clustered relatively closely, while AT/RTs exhibited greater variance across the three groups, with notable outliers such as BT-55 and BT-29. We also performed supervised PCA using all differentially methylated promoter CpG islands (n = 1,058) (**Supplementary Figure S5**). In this analysis, NCH-RBD1 emerged as the only notable outlier, while the remaining MRTs clustered according to tumor classification.

**Figure 4.**
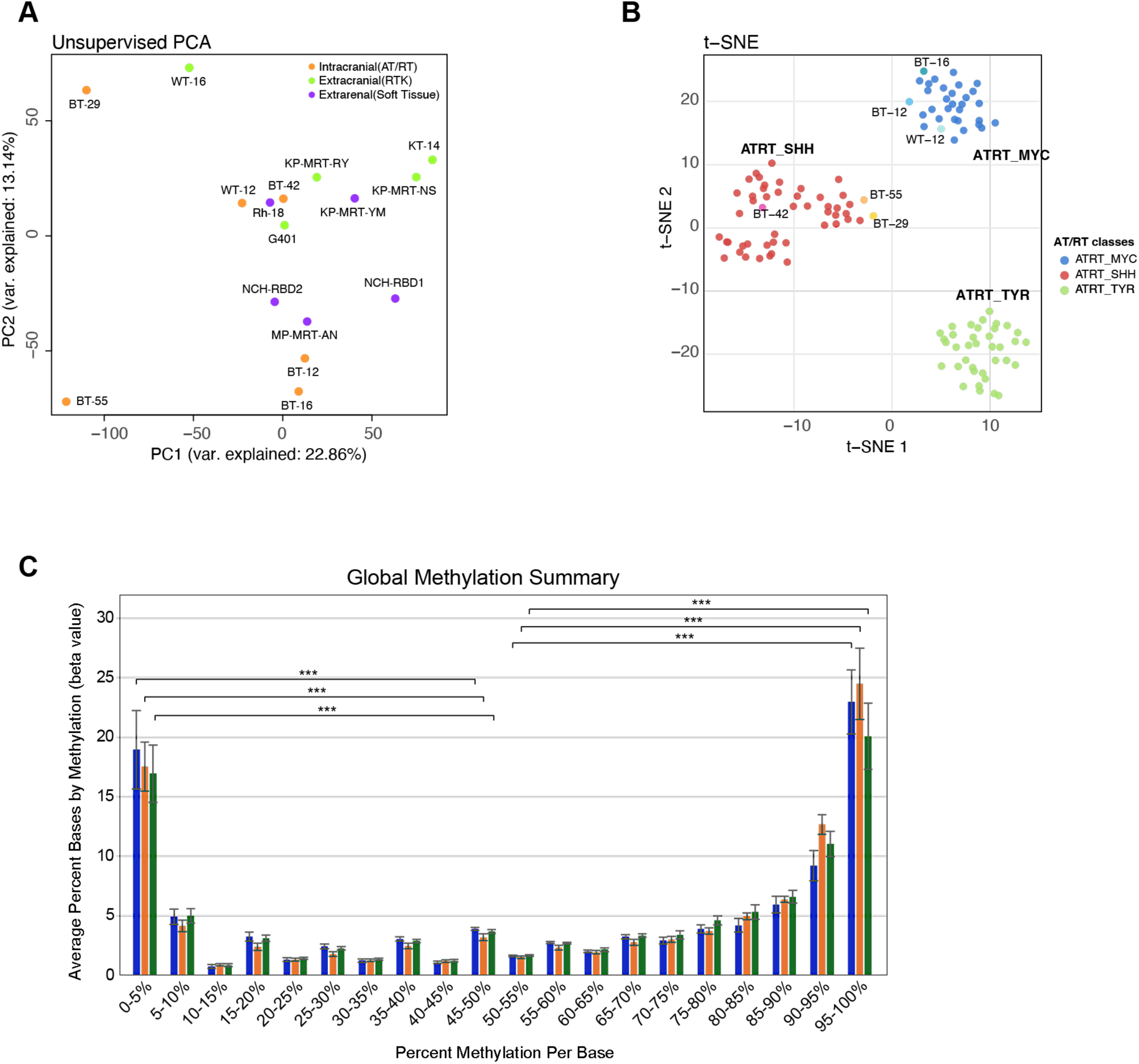
Promoter CpG island methylation analysis of 16 MRT models. (**A**) Unsupervised PCA of all methylation levels of promote CpG islands, colored by respective tumor group. (**B**) t-SNE embedding of genome-wide DNA methylation profiles of six EM-seq AT/RT xenografts shown in the context of AT/RT reference samples from the CNS tumor classification cohort by Capper et al^49^. Reference samples are color-coded by AT/RT subgroup (SHH, MYC, TYR), and EM-seq samples are labeled according to their sample name. (**C**) Global methylation level (®-value) distribution, averaged per tissue group with SEM error bars. T-test p-values are in **Supplementary Table S9**.

Among intracranial tumors, three of the six EM-seq samples were predicted by the MPACT classifier as AT/RT-SHH and clustered together with the AT/RT-SHH cases from the CNS tumor DNA methylation reference cohort described by ^49^, which was restricted to the AT/RT-SHH, AT/RT-MYC, and AT/RT-TYR subgroups. The remaining three EM-seq samples were predicted as AT/RT-MYC and likewise clustered with the AT/RT-MYC reference samples in the t-SNE embedding (**Figure 4B; Supplementary Table S8; Supplementary Figure S4**).

To further dissect methylation patterns, we analyzed β-value distributions (0-100%) as a Global Methylation Summary (**Figure 4C**). Across all groups, methylation polarization was observed. AT/RTs demonstrated a slightly higher β-value concentration in the lowest 0 - 5% range, whereas RTKs had the highest density in the 90 - 100% methylation category. Within each tissue group, differences between extreme and mid-range β-values were significant (**Supplementary Table S9**).

To identify tumor origin-specific methylation patterns, we performed differential methylation analysis focusing on promoter CpG islands. Volcano plots (**Figure 5A, left to right**) illustrate hypo- and hypermethylated promoters for comparisons of AT/RT versus RTK (**Supplementary Table S10**), diagnosis versus relapse (**Supplementary Table S11**), and good versus poor responders (**Supplementary Table S12**). To examine the relationship between DNA methylation and gene expression, **Figure 5B** presents combined analyses for the groups. Only the 2^nd^ and 4^th^ quadrants were considered, as points in these quadrants indicate an inverse relationship between promoter methylation alteration and gene expression^72,73^. Points from DESeq were distributed more evenly in the AT/RT vs RTK comparison, and more abundant in quadrant I for good vs poor responders (**Supplementary Tables S13-15**). Less than 10 genes were significant in both methylation and expression datasets per comparison. The most significant genes in the DNA methylation versus RNA expression correlation plots were *ECHDC2* for AT/RT versus RTK, *ACCS* for diagnosis versus relapse samples, and *B3GALNT1* for good versus poor responders (**Figure 5C**). Volcano and quadrant plots for comparisons involving soft tissue tumors are available in **Supplementary Figure S6**. Significant point information is available in **Supplemental Tables S16-19**.

**Figure 5.**
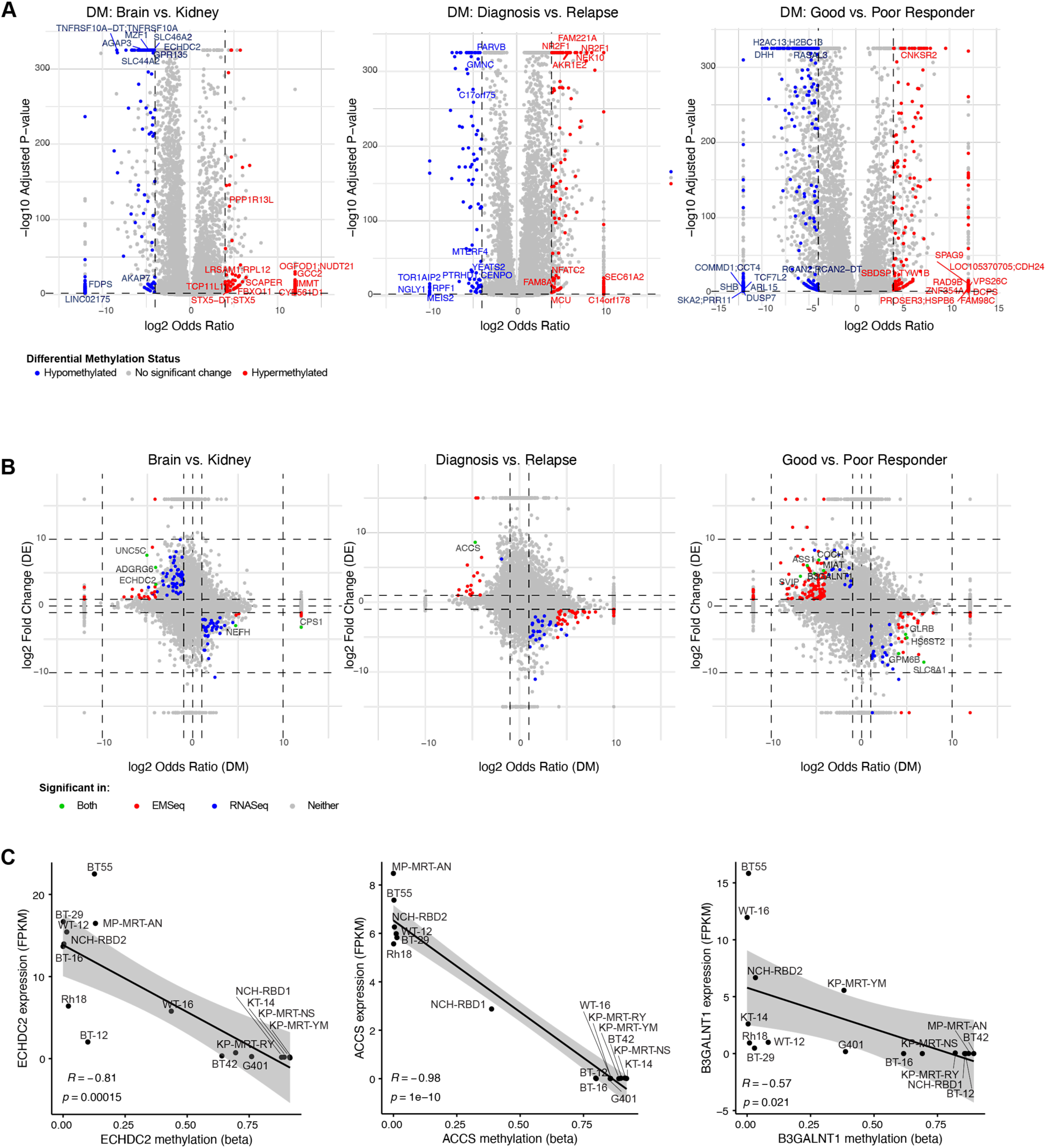
Differential methylation of 16 MRT models across different tissue of origins, tissue collection status, and treatment response. (**A**) Differential methylation volcano plots (from left to right) for Brain vs. Kidney, Diagnosis vs. Relapse, and Good vs. Poor Responder comparisons. Significant differentially methylated promoter CpG islands (colored) were selected based on following criteria: (1) multiple tests adjusted p-value < 0.01; (2) absolute(log2-OR) > 4, and (3) average coverage per feature in both groups > 10 (at least 10x sequencing coverage). Only top 20 hypo- and hyper-methylated promoter CpG islands (their associated gene names) are labeled in these plots. Lists of all DM CpG islands/genes are provided in **Supplementary Tables 6-10**. (**B**) DM versus DE quadrant plots for (from left to right) “Brain vs. Kidney”, “Diagnosis vs. Relapse”, and “Good vs. Poor Responder” comparisons. Genes were significant in DM, in DE, significant in both DM and DE (labeled), or neither, were plotted in red, blue, green and gray, respectively. Lists of all islands/genes are provided in **Supplementary Tables 11-15**. (**C**) Scatter plots of gene expression versus methylation level for genes (*ECHDC2, ACCS*, and *B3GALNT1*) identified as significant in both DM and DE in their respective plots in Figure 5B. Linear regressions and their 95% confidence interval were provided in each plot along with their Pearson correlation coefficients and p-values.

To specifically examine DNA damage repair genes, we selected a panel for closer inspection, even if they did not meet thresholds for differential methylation or expression. Correlation coefficients between methylation and expression are shown in **Figure 6**. Positive correlations were observed for *BRCA1*, *CDK4*, *ATR*, *ABL1*, and *RAD51D*, whereas negative correlations were found for *CDC25A*, *CDK6*, *MYC*, *SLFN11*, and *MGMT* (**Figure 6A**). Notably, *SLFN11* (*Schlafen 11*), known to correlate with chemotherapy sensitivity, particularly to DNA-damaging agents^74,75^, showed a negative correlation with RNA expression (R=0.48, p=0.059) (**Figure 6B**). Similarly, *MGMT* (*O*^6^*-methylguanine methyltransferase*) expression was moderately negatively correlated with methylation (R=0.5, p=0.051) (**Figure 6C**), with β-value visualization near its promoter CpG island (green bar), supporting and expanding our previous findings^25^. We also observed that *LIF* (leukemia inhibitory factor) methylation was strongly negatively correlated with RNA expression (R=0.89, p= 5×10^-6^) (**Figure 6D**). LIF activates multiple signaling pathways, including STAT3, AKT, MAPK, and mTOR, *via* its receptor LIFR^76,77^. The LIF/LIFR axis has been implicated in tumor growth and progression through altering oncogenic processes^76,78^, although its role in MRTs remains uncharacterized.

**Figure 6.**
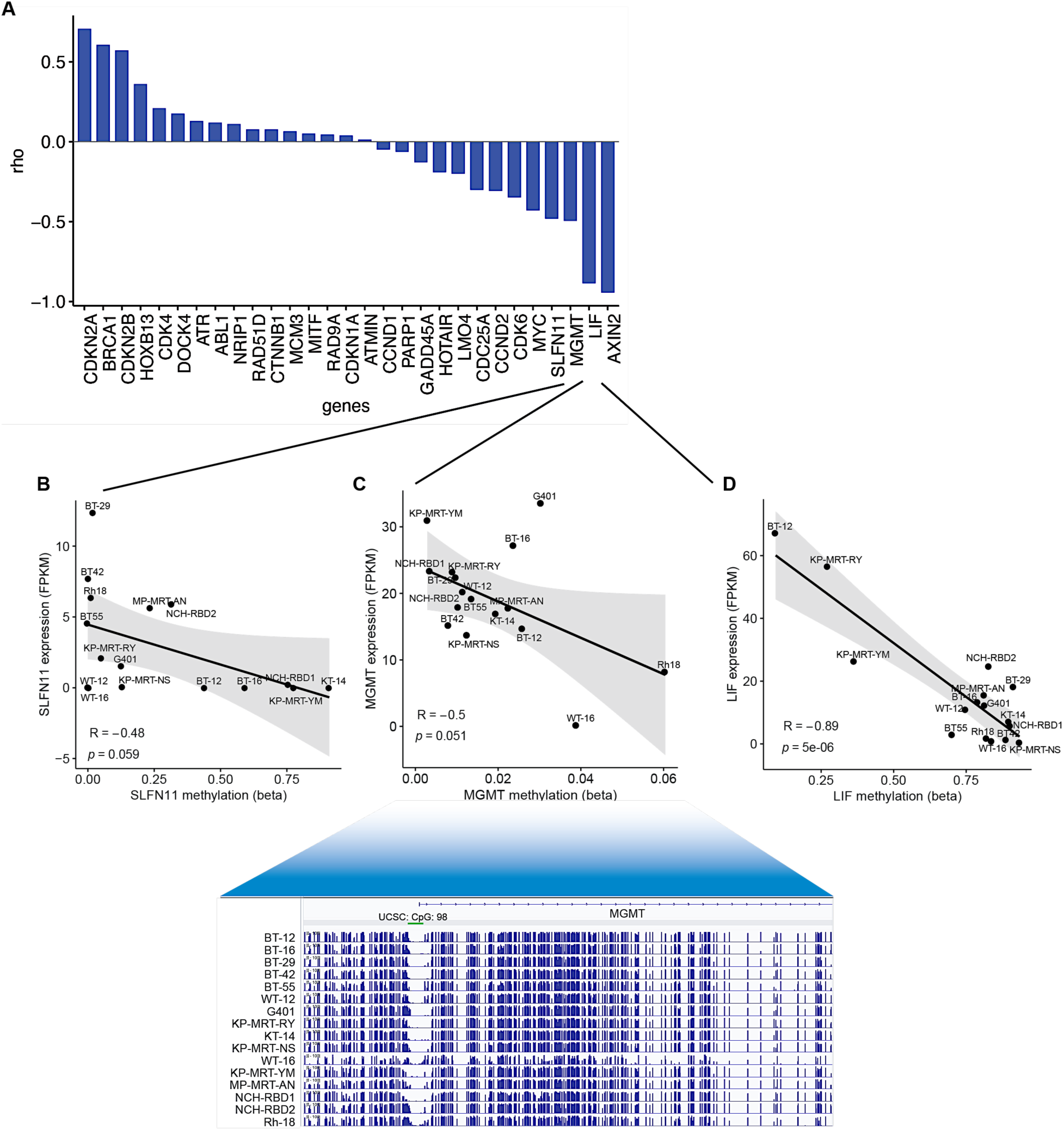
Selected DNA damage-repair genes in MRTs and their association with epigenetic and transcriptional alterations. (**A**) Waterfall plot of a selected panel of DNA damage–repair genes, where the y-axis shows the Pearson correlation between DNA methylation (β-values) and gene expression (FPKM). (**B**) For negatively correlated genes such as *SLFN11*, *MGMT*, and *LIF*, regression plots with sample labels are shown to illustrate methylation-expression relationships across individual MRT samples. For *MGMT*, the genome browser tracks β-values across the promoter region for all 16 MRT samples are shown. USCS CpG island 98, proximal to the *MGMT* transcription start site (TSS), is annotated at the top in green.

## DISCUSSION

Malignant rhabdoid tumors (MRTs) are classically defined by biallelic loss of *SMARCB1* gene (or less commonly *SMARCA4*); however, the mechanisms by which this loss occurs, as well as the extent of additional cooperating alterations remain incompletely understood. In this study, we performed integrated genomic, transcriptomic, and epigenomic analyses of 16 MRT models spanning intracranial, renal, and soft-tissue origins. While MRTs are classically defined by biallelic loss of *SMARCB1*, our data reveal substantial heterogeneity in the mechanisms leading to *SMARCB1* inactivation and uncover broader disruption of genome maintenance pathways that may contribute to tumor pathogenesis, progression, and therapeutic response^3,8,79^.

Only two tumors in our cohort harbored detectable coding single-nucleotide variants (SNVs) in *SMARCB1*. Instead, the predominant mechanisms of *SMARCB1* inactivation involved large-scale deletions and broad loss-of-heterozygosity (LOH) across chromosome 22, strongly supporting allelic loss as a frequent “second hit.” LOH was observed in 5 of 6 intracranial AT/RTs consistent with data reported for low-grade diffusely infiltrated tumors^80^. These findings suggest that deletions and LOH of *SMARCB1* locus represent the dominant mechanisms of *SMARCB1* inactivation in MRTs, rather than recurrent point mutations. This observation reconciles the apparent paradox of preserved coding sequence integrity in many tumors with the near-universal absence of SMARCB1 protein expression, as confirmed by Western blot analysis. In contrast, *SMARCA4* remained intact across all samples, with no evidence of SNVs, indels, copy-number loss, or LOH, indicating the mutual exclusivity of *SMARCB1* and *SMARCA4* alterations in MRTs. Together, these data further refine the genomic definition of MRTs, emphasizing that structural and allelic alterations, rather than point mutations, are the dominant drivers of SWI/SNF complex disruption, reinforcing the concept that *SMARCB1* deficiency is a foundational event in MRT biology, but its genetic basis is more structurally complex than previously appreciated.

Notably, despite the overall low somatic mutation burden characteristic of MRTs, recurrent variants were observed at notable frequency, including *TP53* (P72R, 68.7%) *and BRCA1* single-nucleotide polymorphisms (SNPs; 12.5%). *TP53* missense mutations were detected in 10 of 16 tumors (in 4 of 5 RTKs, 3 of 6 AT/RTs, and 3 of 5 of soft tissue MRTs), all carrying the P72R variant, with one RTK sample (KP-MRT-NS) harboring an additional more pathogenic R273C mutation^81^. While the P72R variant retains partial tumor suppressor function^59^, its co-occurrence with alterations in *BRCA1* and *BRCA2*, detected in 2 and 3 of 16 MRTs, respectively, including one relapse sample (WT-16) with mutations in both genes, suggests a broader perturbation of DNA damage response (DDR) pathways in a substantial subset of tumors. As PDX development is not known to introduce *BRCA1/2* mutations^82^, these alterations may reflect tumor-intrinsic genomic effects. Although most identified *BRCA1* or *BRCA2* SNPs are classified as likely benign, their enrichment in *TP53*-altered tumors, together with the presence of mutational signature SBS5, strengthens the evidence for underlying defects in genome maintenance. SBS5 has previously been linked to defects in nucleotide excision repair and increased genomic instability resulting from unrepaired DNA damage^83^. Taken together, these findings support a model in which *SMARCB1* loss occurs within a genomic context already compromised in DNA integrity, thereby facilitating tumor initiation and progression despite the characteristically low somatic mutation burden associated with MRTs^84^.

Although MRTs are often described as genomically simple, both prior studies and our whole-genome analysis revealed extensive structural variation (SV) across the cohort, with an average of over 475 SVs per tumor^85^. While most of SVs were deletions, selected samples exhibited marked chromosomal rearrangements and focal copy-number gains^86^. Specifically, partial duplications of *AHI1* and *MYB* in the KP-MRT-NS model, together with the validated *AHI1:MYB* fusion transcript, point to additional oncogenic mechanisms in a subset of MRTs. However, despite the use of stringent selection criteria, these structural variants and gene fusions remain susceptible to technical bias and alignment artifacts, and therefore warrant further validation, ideally in matched primary patient samples. In addition, the functional consequences of the *AHI1:MYB* fusion have yet to be defined. Nevertheless, *MYB* dysregulation is well established in other pediatric malignancies, suggesting that this fusion may contribute to transcriptional reprogramming. For instance, *MYB:QKI* fusions have recently been reported in pediatric-type diffuse low-grade glioma, supporting a broader oncogenic role for *MYB* rearrangements^87^. Notably, the KP-MRT-NS model harboring the unique *AHI1:MYB* fusion was derived from a previously treated patient (therapy unknown) and also carries an oncogenic *TP53* mutation (R273C). These features may reflect molecular changes acquired during treatment or relapse evolution and provide insights into clonal dynamics and potential biomarkers of disease progression.

Although MRTs share common genetic hallmarks, they showed pronounced transcriptomic heterogeneity driven by tissue of origin, disease status, and therapeutic response^7^. Differential expression analyses revealed biologically coherent, subtype-specific programs, including interferon signaling in AT/RTs, xenobiotic metabolism in kidney MRTs, and epithelial-mesenchymal transition in soft-tissue tumors^88^. These data suggest that cell-of-origin and microenvironmental context significantly shape MRT biology beyond core genetic drivers. Relapse-associated MRTs displayed transcriptional changes consistent with upregulation of *embryonal Fyn-associated substrate* (*EFS)* and *fibroblast growth factor 19 (FGF19)* and downregulation of *calcium binding protein (S100A11)* and *protein kinase C delta binding protein* (*PRKCDBP)* genes*. PRKCDBP* and *S100A11* are broadly described as context-dependent tumor suppressors, while *EFS* and *FGF19* have been implicated in oncogenic signaling and migration and adhesion, respectively^89–92^. While the small number of relapse samples limits definitive conclusions, these findings suggest adaptive transcriptional remodeling during disease progression or treatment pressure. Importantly, broad transcriptomic profiling also distinguished “good” *versus* “poor” MRT responders to PEGylated PARP inhibitor talazoparib and DNA alkylating agent temozolomide^25^. Enrichment of epidermal growth factor receptor (EGFR), Ephrin family members, and receptor tyrosine kinase (RTK) signaling in “good” responders indicates that heightened proliferative or DNA damage-associated signaling states may sensitize tumors to PARP inhibition. Consistent with this model, elevated EGFR expression observed in responding models as well as in untreated tumors, suggests its potential role as a predictive biomarker of therapeutic vulnerability.

DNA methylation is a key epigenetic regulator of cell identity and gene expression, with tissue-and context-specific patterns that influence development, disease states, and environmental responses. The 6 AT/RT samples segregated into SHH and MYC subgroups, consistent with the established methylation and transcriptomic-based classifiers, validating model fidelity^56^. Differential methylation analyses identified promoter CpG island changes associated with tissue of origin, disease progression, and therapeutic response. Unsupervised and supervised analyses demonstrated that kidney and soft-tissue MRTs (including 2 samples derived from liver, one from chest-wall, and one from perineal tissue) formed a relatively tight cluster, whereas AT/RTs displayed greater epigenetic variability, indicating increased plasticity or heterogeneity within intracranial tumors. Notably, NCH-RBD1, the only soft tissue MRT derived from lung tissue, segregated from all other soft tissue MRTs, consistent with a distinct epigenetic state.

Integration of methylation and expression data revealed relatively limited overlap, indicating that while promoter methylation contributes to transcriptional regulation in MRTs, additional mechanisms, including chromatin remodeling and transcription factor activity, are likely dominant drivers^93^. Notably, focused analysis of DNA repair genes revealed significant correlations between promoter methylation and gene expression, highlighting one of the well-known mechanisms of gene silencing through promoter hypermethylation. Prior evidence indicates that hypermethylation-associated suppression of *SLFN11* (*Schlafen 11*) and *MGMT* (*O*^6^*-methylguanine methyltransferase*) gene expression is linked to chemotherapy sensitivity^26,94–100^. The inverse correlation observed between *SLFN11* promoter methylation and gene expression is especially intriguing, as *SLFN11* has been shown to modulate response to DNA-damaging agents and PARP inhibitors. Similarly, *MGMT* methylation is a well-established determinant of glioblastoma response to alkylating agents such as temozolomide, reinforcing the translational importance of epigenetic profiling of this gene. Our analysis revealed that the two MRT models with the most favorable responses to PARP inhibition and DNA damaging therapy (WT-16 and Rh-18)^25^, also had the highest levels of *MGMT* promoter methylation. Furthermore, a strong inverse relationship between promoter methylation and expression of the immune modulator *LIF* (*leukemia inhibitory factor*) points to the *LIF/LIFR* (*LIF receptor*) axis as a potentially important, yet previously underexplored, pathway in MRT biology^101^. *LIF* is highly expressed in MRTs, and its expression is inversely correlated with SWI/SNF complex activity. Given *LIF*’s known role in activating STAT3, AKT, MAPK, and mTOR signaling pathways, dysregulation of this axis may contribute to tumor progression or therapeutic resistance and merits further functional investigation^102,103^.

By integrating genomic, transcriptomic, and epigenomic data, we show that MRTs, while defined by *SMARCB1* loss, harbor substantial structural and regulatory heterogeneity. These findings expand the molecular landscape of MRTs and identify biomarkers and pathways with potential to guide the development of rational, precision treatments.

## STUDY LIMITATIONS

Given the extreme rarity of MRTs, we assembled a substantial collection of MRT xenograft models capturing distinct genomic, transcriptomic, and epigenetic programs, providing a valuable resource for future biological investigation. Nevertheless, this study has unavoidable limitations related to cohort size and reliance on xenograft models, which may not fully recapitulate tumor-microenvironment interactions. Matched patient germline DNA was unavailable, largely due to historically limited access to non-tumor tissue from patients and the retrospective nature of sample collection. While germline data would have enabled more accurate discrimination of somatic from inherited variants, we mitigated this limitation through stringent population-based filtering and cross-model comparisons. Additionally, the EMseq platform used here primarily targets promoter-associated CpG islands, limiting the breadth of epigenomic coverage. While this conventional approach facilitates integration with transcriptomic data, a substantial proportion of regulatory epigenetic information remains unexplored. Despite these constraints, our dataset represents one of the most comprehensive molecular characterizations of MRT xenografts to date and provides a foundation for mechanistic and therapeutic studies.

## ACKNOWLEDGEMENTS

The authors gratefully acknowledge the patients who consented to the donation of tissues and cells used to generate the experimental models that enabled preclinical investigation of novel therapeutic agents and multi-omic analyses.

